# Tailoring a CRISPR/Cas-based Epi-editor for Decreased Cytotoxicity and Enzymatic Specificity

**DOI:** 10.1101/2024.09.22.611000

**Authors:** Jacob Goell, Ruopu Jiao, Jing Li, Alex J. Ma, Kevin Gong, Guy C. Bedford, Daniel Reed, Barun Mahata, Sunghwan Kim, Spencer Shah, Shriya Shah, Maria Contreras, Suchir Misra, Keith E. Maier, Michael R. Diehl, Mario Escobar, Isaac B. Hilton

## Abstract

CRISPR-based epi-editors can robustly modulate cellular transcription and chromatin structure, but off-target activity and cytotoxicity limit their utility. Here, we engineer the acyl-CoA binding pocket of the human p300 histone acyltransferase to reduce its cytotoxicity and tune its acylation profile when fused to dCas9. We discover a single amino acid substitution (I1417N) that decreases cytotoxicity related to exogenous p300 overexpression yet preserves dCas9-p300 mediated histone acetylation and gene activation. We find that dCas9-p300 I1417N is less perturbative to the transcriptome and proteome of human cells, and that this behavior is driven by favorable stability kinetics. We also develop a crotonylation-biased dCas-p300 variant (I1395G) that selectively deposits histone crotonylation and activates transcription from endogenous promoters at levels comparable to wild-type p300. The p300 variants generated here enhance epi-editing capabilities and demonstrate that engineering of catalytic domains can be a powerful strategy for tailoring enzymatic activities and mitigating effector-driven toxicity in epi-editing.

## Main

Epigenome editing (epi-editing) enables the modulation of human gene expression and chromatin structure for the study of epigenomic regulatory mechanisms and the development of new therapeutics^1–4^. Epi-editing platforms leverage DNA-binding domains, such as deactivated CRISPR-Cas (dCas) proteins, as a scaffold for bringing epigenetic effectors to targeted genomic loci. Effectors are often repurposed from viral or eukaryotic gene-regulatory proteins and function to recruit transcriptional machinery or cofactors, change the methylation state of DNA, or catalyze histone post-translational modifications (PTMs)^5^. This approach has enabled the development of epi-editing tools that have furthered our understanding of heritable gene silencing^6,7^, enhancer biology^8–11^, and synthetic transcriptional activation and repression^12–16^. Despite this promise, many of the most potent and widely-used epi-editors that directly catalyze chemical modifications to histones or DNA are limited by pervasive toxicity upon exogenous expression^6,17–20^.

Previously, we engineered a potent transcriptional activation and histone lysine acylation tool by fusing the catalytic core domain of the human EP300 (p300) protein to dCas9 (dCas9-p300)^21^. The dCas9-p300 fusion directly catalyzes histone lysine acetylation at targeted genomic loci and is able to activate transcription form both enhancer and promoter elements. The p300 protein is of particular interest as an epi-editing effector as p300 is a central regulator of histone lysine acetylation, enhancer activity, and signal-dependent gene activation in vertebrates^22–24^. Further, dysregulation of p300 is linked to numerous human diseases, including diverse cancers and neurodevelopmental disorders^25,26^. While the ability to leverage dCas9-p300 for epi-editing is extremely powerful, overexpression of p300 or the dCas9-p300 fusion has been linked to inhibited cell growth, cytotoxicity, and cellular senescence^11,20,22,27,28^. As such, mitigating the cytotoxicity and off-target effects of dCas9-p300 would enable new opportunities for mechanistic epigenetics and cell engineering, as well as for disease modeling and focused therapeutic interventions.

In addition to acetylation, the acyltransferase core of p300 can also catalyze longer-chain acylations such as propionyl, butyryl, crotonyl, and lactyl^21,29,30^. Histone lysine acylation is a dynamic and highly regulated PTM that guides appropriate gene expression in response to various stimuli^31–38^. As the substrates for these longer chain acylations are derived through different metabolic processes, these PTMs are highly context-specific and are often only present in cell types wherein specific metabolic pathways are active and corresponding feedstocks are available. Longer-chain acylations are chemically distinct from acetylation and exhibit different affinities for epigenetic reader proteins^39–41^, however the study of these marks has been relatively underexplored, due in part to the lack of tools to interrogate the roles that different acylations play at endogenous loci.

Mutagenesis of the p300 Acyl-CoA domain has previously been shown to alter the enzymes substrate preference or modulate acylation activity to varying degrees^21,30,42,43^. However, to our knowledge, this technique has not yet been explored for reducing p300 cytotoxicity. Here, we used targeted mutagenesis of the human p300 Acyl-CoA binding domain to discover and benchmark a p300 variant (I1417N) that substantially reduced cytotoxicity and off-target perturbations associated with exogenous expression of the wild-type p300 core (p300 WT), without sacrificing the ability to deposit lysine acetylation and activate transcription. We found that the reduction in perturbative behavior caused by the I1417N mutation is likely due to destabilization of the protein into a pseudo-stable state wherein on-target activity is possible, but the protein is degraded before off-targets can be incurred, a strategy that could likely be applied to other epi-editing effectors.

We additionally engineered a programmable lysine crotonyltransferase by incorporating a recently described mutation (I1395G) that skews catalytic activity toward crotonylation into dCas9-p300, and leveraged this tool to gain insights into the divergent roles that histone acylation marks can have within the human genome. Altogether, we envision that the engineering strategies and tools we have developed will be instructive for understanding the role of lysine acylation in transcriptional control and for improving epi-editing tools beyond those applied for programmable histone acylation.

## Results

### Targeted mutagenesis of the p300 core domain generates functional variants with altered transactivation potential and expression levels

Given that single amino acid substitutions in the p300 Acyl-CoA binding region have previously been observed to alter the catalytic behavior of p300^21,30,42,43^, we hypothesized that other mutations within the p300 Acyl-CoA binding region could result in variants with reduced cytotoxicity through modulation of p300 catalytic activity. We first identified 9 different amino acid substitutions within the p300 Acyl-CoA binding region that do not occur in the Catalogue of Somatic Mutations in Cancer (COSMIC)^44^ or ClinVar^45^ databases (**Fig. 1a, Extended Data Fig. 1a**). We next cloned p300 variants bearing these mutations into reverse tetracycline repressor (rTetR) fusion constructs, along with p300 WT, the catalytically inactivated p300 D1399Y, and p300 I1395G, a missense mutation previously characterized to bias the catalytic activity of p300 towards lysine crotonylation^43^. These rTetR-p300 variants were bicistronically expressed with mCherry through linkage via a porcine teschovirus self-cleaving peptide (P2A), decoupling the protein stability of each variant from mCherry expression. To screen for activation potential, we engineered an AAVS1 knock-in HEK293T reporter cell line with 9x TetO binding sites adjacent to a minimal SV40 promoter, driving the expression of a fluorescent Citrine reporter when activated^46^ (**Fig. 1b**).

**Fig 1:**
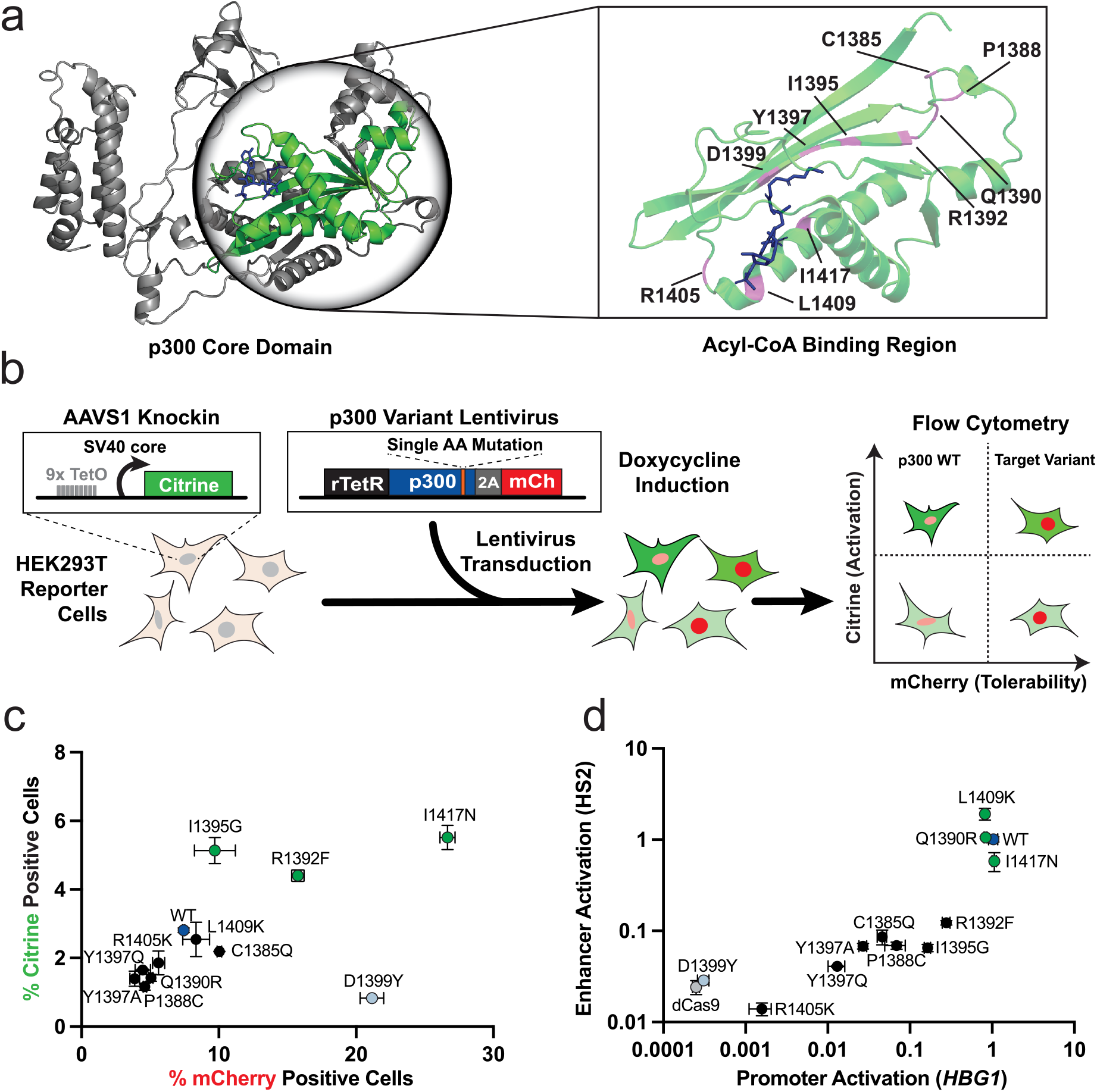
Targeted mutagenesis of the p300 acyl-CoA binding region produces variants with altered expression and transactivation potential. **a**, Crystal structure of the p300 core domain (5LKU) with the Acyl-CoA binding region colored in green and Acyl-CoA molecule in blue. Expanded view of the Acyl-CoA binding region displaying the residues targeted for mutation in this study highlighted in magenta. **b**, A schematic depicting the rTetR-rTetO screening system. 9x TetO binding sites adjacent to a minimal SV40 promoter immediately upstream of a fluorescent Citrine reporter were knocked-in to the *AAVS1* safe harbor locus. Engineered HEK293T cells were transduced with indicated variants bicistronically expressed with mCherry. Doxycycline mediated recruitment of functional rTetR-p300 variants to the TetO binding sites activated Citrine expression. Citrine and mCherry fluorescence were read out via flow-cytometry, with Citrine % positivity serving as a readout for activation potential and mCherry % positivity serving as a readout for relative expression. **c**, mCherry-Citrine flow cytometry percent positivity for the reporter line cells transduced with the tested p300 variants, collected 48 hours post-doxycycline addition. Variants that outperformed p300 WT are colored in green (n = 2, mean ± SEM). **d**, RT-qPCR for *HBG1* transcriptional activation 72 hours after transient transfection of the tested dCas9-p300 variants with 4 gRNAs targeted to the HS2 enhancer (Y-axis) or 4 gRNAs targeted to the *HBG1* promoter (X-axis) in HEK293T cells (n = 3-8, mean ± SEM).

Transduction with rTetR-p300 WT and rTetR-p300 D1399Y demonstrated expected trends of gene activation and histone acetylation in the presence of doxycycline (Dox; **Extended Data Figs. 1b,c**). Cells transduced with the catalytically inactivated rTetR-p300 D1399Y expressed the bicistronic mCherry transgene to a greater degree than rTetR-p300 WT transduced cells, demonstrating that p300 WT induced toxicity impacted the ability of cells to express mCherry - enabling us to use mCherry fluorescence as a readout for cell tolerance of p300 variants. Direct fusion of mCherry to the N-terminus of dCas9-p300 WT or D1399Y recapitulated the differences in mCherry expression observed with the bicistronic system (**Extended Data Figs. 1d,e**).

We next assessed the activation potential (Citrine output) and expression tolerability (mCherry level) for the full panel of selected p300 variants. All tested variants displayed reporter activation levels that exceeded the catalytically dead p300 D1399Y. Interestingly, three variants (I1417N, I1395G, and R1392F) outperformed rTetR-p300 WT with respect to both reporter gene activation and mCherry expression (**Fig. 1c**). We then fused each p300 variant to the C-terminus of dCas9 to assess relative activation potential at endogenous loci, using the HS2 enhancer and cognate *HBG1* promoter as testbeds. We observed that three of the engineered variants (I1417N, Q1390R, and L1409K) performed comparably to p300 WT when targeted to either the HS2 enhancer or *HBG1* promoter using corresponding gRNAs (**Fig. 1d**). While many of the tested variants only weakly activated Citrine expression compared to the D1399Y variant in the rTetR-TetO system, most activated the endogenous *HBG1* gene to a much greater relative degree when fused to dCas9, highlighting the importance of both chromatin context and recruitment architecture in the design and testing of epi-editors.

A negative correlation between activation potential and cell viability has previously been observed with other epi-editing activators^11^. Therefore, we assessed whether this trend was generalizable to variants of dCas9-p300. We screened a library of 800 substitution mutations and 40 single amino acid deletions across a contiguous 40 amino acid stretch of the Acyl-CoA binding domain of p300 and indeed observed a negative correlation between activation potential and cell viability (**Extended Data Fig. 1f, Supplementary Table 1**). Collectively, these data indicate that the cytotoxicity associated with exogeneous expression of dCas9-p300 WT is linked to the mechanisms by which dCas9-p300 activates transcription, and that this relationship can be modulated through mutagenesis of the Acyl-CoA binding pocket.

### dCas9-p300 I1417N displays improved expression in human cells without sacrificing histone acetylation nor gene activation potential

p300 I1417N was the only variant that outperformed p300 WT in gene activation and expression tolerability in the rTetR-rTetO system and that was comparable to p300 WT at activating transcription when targeted to either the *HBG1* promoter or the HS2 enhancer (**Fig. 1c,d**). Interestingly, in HEK293T cells transfected with bicistronic mCherry constructs co-expressed with our p300 variants, dCas9-p300 WT displayed less mCherry fluorescence than dCas9-p300 I1417N, which exhibited mCherry levels that were similar to dCas9 alone and dCas9-p300 D1399Y (**Fig. 2a**). This finding was recapitulated using dCas9-p300 variants directly fused to the N-terminus of mCherry (**Extended data Fig. 2a**). We also directly assessed the protein expression levels of the dCas9-p300 variants 72 hours after plasmid transfection into HEK293T cells and found that dCas9-p300 I1417N was more abundantly expressed than dCas9-p300 WT (**Extended data Fig. 2b**). We reasoned that these observations were likely due to dCas9-p300 I1417N expression resulting in less cytotoxicity when compared to dCas9-p300 WT. Indeed, using alamarBlue to readout cell count from primary donor-derived Mesenchymal Stem Cells (MSCs), we found that in these relatively sensitive cells expression of dCas9-p300 I1417N resulted in no significant loss of cell count when compared to dCas9, while expression of dCas9-p300 WT resulted in substantial toxicity (**Fig. 2b**). Similar effects were observed in K562 cells, a workhorse cell line for epi-editing screens and experimentation^9,11,47^ **(Extended data Fig. 2c)**. In addition to decreased cytotoxicity, we found that when packaging dCas9-p300 variants in lentiviral vectors and quantifying their physical titers, dCas9-p300 I1417N had a 2-fold greater titer than dCas9-p300 WT, as well as a 5-fold improvement in the percentage of transduced K562 cells expressing the lentivirus cargo (**Extended data Fig. 2d**). These data demonstrate that the single I1417N mutation in p300 mitigates cytotoxicity in human cells which can manifest as improved construct expression in plasmid and lentiviral-based experimental contexts.

**Fig 2:**
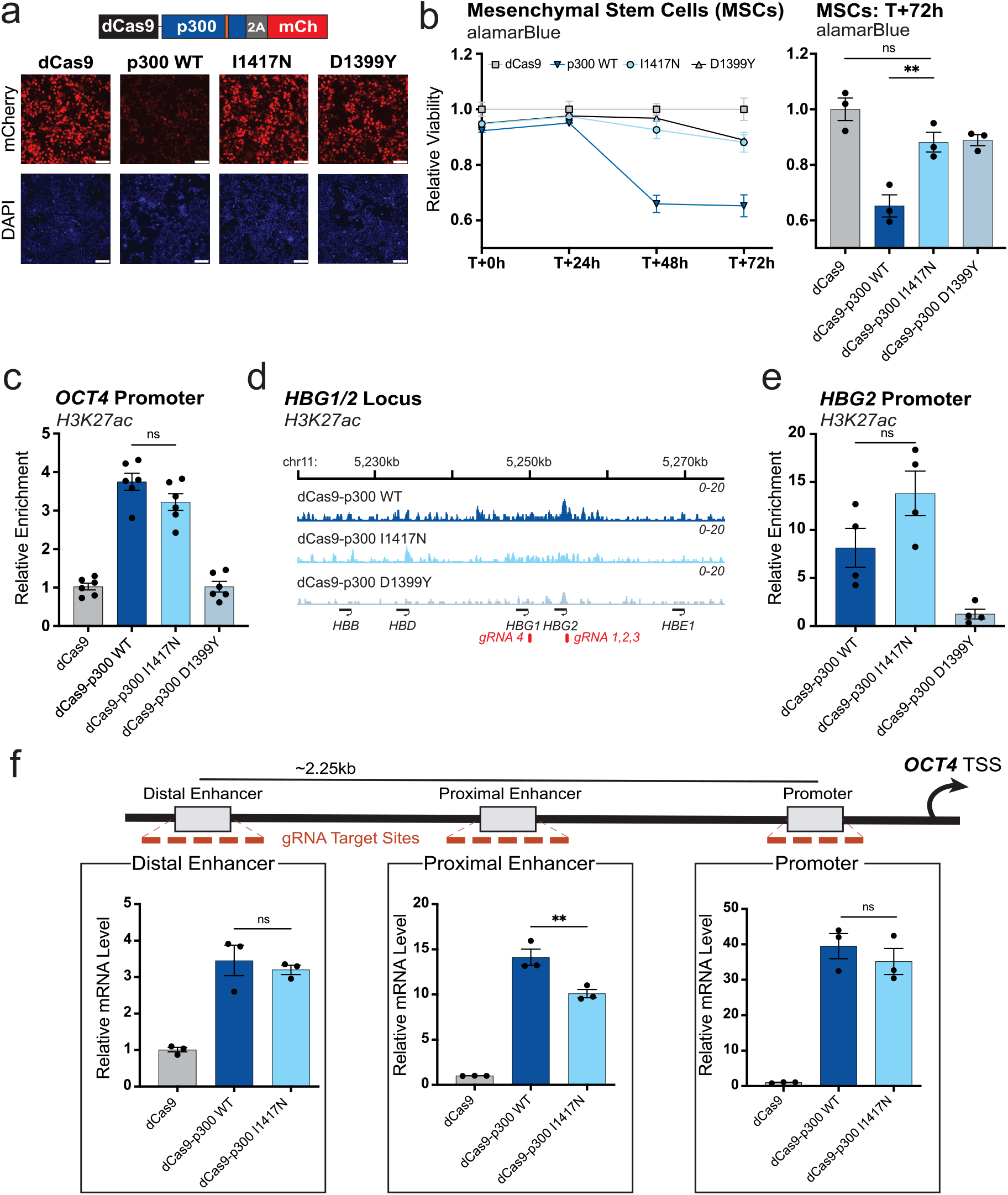
p300 I1417N deposits histone acetylation and activates gene expression from endogenous regulatory elements without cytotoxicity. **a**, Fluorescence microscopy 48 hours after HEK293T cells were transiently transfected with dCas9, dCas9-p300 WT, dCas9-p300 I1417N, or dCas9-p300, all bicistronically expressed with mCherry and co-transfected with a scrambled gRNA. Top panels: mCherry channel fluorescence. Bottom panels: DAPI staining. Scale bar: 100um. Orange line in schematic indicates approximate location of p300 variant mutation. **b**, Relative cell viability of primary donor-derived Mesenchymal Stem Cells (MSCs) after transfection with mRNA constructs encoding either dCas9 alone, or dCas9 fused to the indicated p300 variants. Left panel: Relative cell viability assessed via AlamarBlue relative to the dCas9 treated MSCs at 0, 24, 48, or 72 hours after transfection. Right panel: Relative MSC viability 72 hours post transfection with indicated variants (n = 3, mean ± SEM). **c**, CUT&RUN qPCR for H3K27ac at the *OCT4* promoter in HEK293T cells 60 hours after transient transfection with the indicated constructs and sgRNAs targeting the *OCT4* promoter. Relative enrichment was calculated as fold change over dCas9 (n = 6, mean ± SEM). **d**, H3K27ac CUT&RUN sequencing reads from HEK293T cells collected after transient transfection with the indicated dCas9-p300 variant and 4 *HBG1-*targeting sgRNAs are shown mapped for genomic coordinates spanning ∼5,220,000 to ∼ 5,275,000 bp of human chromosome 11 (GRCh38/hg38, n=2). Relevant gene annotations and locations of gRNAs and CUT&RUN qPCR primers are shown in the bottom row. **e**, CUT&RUN qPCR for H3K27ac at the *HBG2* promoter in HEK293T cells 60 hours after transient transfection with the indicated constructs and gRNAs (n = 4, mean ± SEM). **f**, Top: Schematic of the *OCT4* locus with enhancer and promoter elements arranged relative to the transcription start site (TSS). Red rectangles indicate target gRNA locations. Bottom: RT-qPCR for *OCT4* transcript expression in HEK293T cells 72 hours after plasmid transfection with the indicated constructs and sgRNAs targeting the *OCT4* distal enhancer, proximal enhancer, and promoter (n = 3, mean ± SEM). Comparison lines throughout indicate multiple comparison testing with Šidák, Dunnett’s T3, or Dunn’s multiple comparisons tests depending on normality, tested via Shapiro-Wilk (*P>0*.*05*), and homoscedasticity, tested via Brown-Forsythe (*P>0*.*05*). ^*^*P*≤*0*.*05*, ^**^*P*≤*0*.*01*, ^***^*P*≤*0*.*001*, ^****^*P*≤*0*.*0001*, ns, not significant.

Given the observed inverse relationship between activation potential and cytotoxicity among dCas9-p300 variants (**Extended data Fig. 1g**), we next assessed the catalytic and transcriptional activity of dCas9-p300 I1417N. Remarkably, we found that targeting the *OCT4* promoter with dCas9-p300 WT and dCas9-p300 I1417N resulted in similar levels of H3K27ac deposition as assayed via CUT&RUN qPCR (**Fig. 2c**). We also found similar levels and patterning of H3K27ac deposition between dCas9-p300 WT and dCas9-p300 I1417N when targeted to the *HBG1/2* locus using CUT&RUN sequencing and CUT&RUN qPCR (**Fig. 2d,e**). Additionally, we performed CUT&RUN qPCR for H2BK20ac, H3K4me3, and H3K27me3 at the *HBG1/2* locus after targeting with dCas9-p300 variants (**Extended data Fig. 2e**). H2BK20ac, a p300/CBP-specific histone modification thought to designate active enhancers^48^, was enriched at similar levels by both dCas9-p300 WT and dCas9-p300 I1417N. H3K4me3, a histone modification shown to have substantial crosstalk with H3K27ac^15,49,50^, was also similarly enriched by both dCas9-p300 WT and dCas9-p300 I1417N. H3K27me3, a polycomb-mediated histone modification associated with repressive chromatin states^51^, was not enriched by either dCas9-p300 WT or dCas9-p300 I1417N.

Concurrent with histone lysine acetylation, transcriptional activation is a hallmark response when dCas9-p300 WT is targeted to gene enhancers and promoters^21^. Targeted transcriptional activation of *OCT4* from either the distal enhancer or promoter was comparable between dCas9-p300 WT and I1417N, while dCas9-p300 I1417N was slightly less effective at the proximal enhancer (**Fig. 2f**). Transcriptional activation with dCas9-p300 I1417N was also comparable to dCas9-p300 WT when targeted to other loci and transfected in different cell lines (**Extended data Fig. 2f,g**). We further observed that p300 I1417N was compatible with orthogonal dCas species and achieved potent gene activation in these configurations (**Extended data Fig. 2h**). Additionally, using a library of perturb-seq gRNAs targeting distal regulatory elements^47^ (**Supplementary data Fig. 1**), we found that the gene activation response profiles for dCas9-p300 WT and dCas9-p300 I1417N were similar in regards to responsive regulatory element distance from cognate transcriptional start sites and the average fold-change observed across hundreds of perturbations (**Extended data Fig. 3a,b**). Intriguingly, when directly comparing the perturbation profile of these dCas9-p300 activators against that of dCas9-VP64/VPR^47^, we found that dCas9-p300 targeting resulted in 43 unique perturbation pairs that were not seen with dCas9-VP64/VPR, with 11 being enhancer element mediated perturbations (**Extended data Fig 3c, Supplementary Table 2**).

Altogether, our findings indicate that although dCas9-p300 I1417N is significantly less cytotoxic than dCas9-p300 WT, no substantial deficits in programmable transcriptional activation or acetylation activity are incurred. This unique feature of dCas9-p300 I1417N establishes this variant as a powerful alternative to dCas9-p300 WT, especially for use in studies in sensitive cells lines or for the development of epi-editing therapeutics where cytotoxicity cannot be tolerated. Finally, despite our finding that increasing activation potential is negatively correlated with cell viability (**Extended Data Fig. 1**), dCas9-p300 I1417N is evidence that outliers from this general trend can be engineered.

### dCas9-p300 I1417N is less perturbative to human cells than dCas9-p300 WT

Many epi-editing technologies display cytotoxicity when exogenously overexpressed, which motivated us to further study the mechanistic basis behind the reduced cytotoxicity seen with dCas9-p300 I1417N compared to dCas9-p300 WT^20^. We hypothesized that the reduced cytotoxicity seen with p300 I1417N was related to protein-protein interactions (PPIs) and/or differential expression kinetics. To capture the proteomic interactions of dCas9-p300 variants, we performed immunoprecipitation followed by mass-spectrometry (IP-MS) on protein gathered from HEK293T cells transiently transfected with dCas9-p300 variants and a scrambled gRNA. We filtered out hits that were highly prevalent in the CRAPome (>50% occurrence) to remove potential false positive interactors that are common in IP-MS experiments^52^, and identified significant differential PPIs for all tested dCas9-p300 variants against a dCas9 only control “bait”. We found that dCas9-p300 WT exhibited the highest number of significant differential interactors (p_val_<0.05) with 253 hits, whereas dCas9-p300 I1417N had far fewer (167 hits) and the catalytically inactivated dCas9-p300 D1399Y mutant exhibited the fewest (117 hits; **Extended Data Fig. 4a, Supplementary Table 3**). Several identified p300 interactors were cross-validated using the STRING database^53^, with p300 WT again capturing the most established interactors, followed by I1417N, and D1399Y (**Extended Data Fig. 4b**). Interestingly, we found that only catalytically active p300 variants associated with Sirtuin 2 (SIRT2), a deacetylase known to interplay closely with p300 in balancing endogenous protein acetylation^54^ (**Extended Data Fig. 4b, c**).

Endogenous human p300 is involved in the acetylation of thousands of proteins across the proteome^22,23,55^. The reduced PPIs seen with dCas9-p300 I1417N suggested that fewer proteins in the proteome were being aberrantly acetylated by dCas9-p300 I1417N. To interrogate this possibility, we compared our IP-MS results with a human p300 acetylome dataset^23^. We considered differentially enriched PPIs found in our IP-MS experiment that were also found in the p300 acetylome to be indicative of a possible aberrant acetylation event. Remarkably, we found that exogenous overexpression of dCas9-p300 WT resulted in over three times as many potential unique aberrant acetylation events when compared to dCas9-p300 I1417N, consistent with the global increases in acetylated lysine residues observed with dCas9-p300 WT but not dCas9-p300 I1417N reported while our study was in preprint^20^ (**Fig. 3a, Supplementary Table 4**).

**Fig 3:**
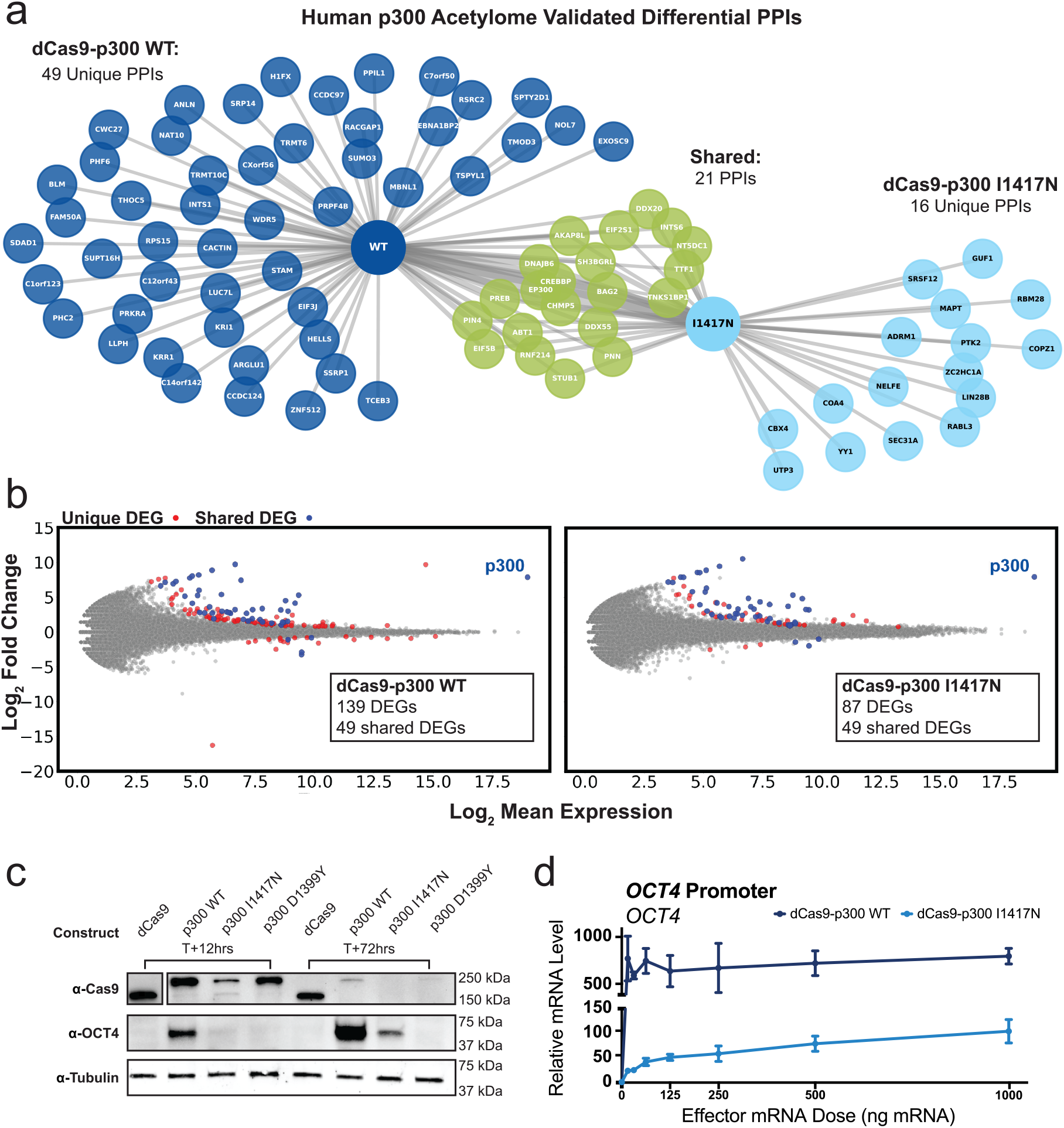
dCas9-p300 I1417N is less perturbative to human cells than dCas9-p300 WT. **a**, Network graph illustrating differential protein-protein interactions (PPIs) found for dCas9-p300 WT and dCas9-p300 I1417N compared to a dCas9 only “bait” based upon the endogenous p300 acetylome. Edges in the graph were distance weighted by the significance of the differential PPI for the p300 variant each protein was connected to. **b**, (Y-axis) Log_2_(Fold Change) for transcripts detected via RNAseq in HEK293T cells transiently transfected with 4 gRNAs targeting the HS2 enhancer element graphed against (X-axis) the Log_2_(Mean Expression) of each transcript for (left) dCas9-p300 WT and (right) dCas9-p300 I1417N, both compared to dCas9 alone. Grey dots indicate transcripts that were not significantly different between the p300 variant and dCas9 (Benjamini-Hochberg corrected *Padj <* 0.05). Red dots indicate significant differentially expressed genes (DEGs) that were unique to the specific p300 variant. Blue dots indicate DEGs that were shared between dCas9-p300 WT and dCas9-p300 I1417N (n = 2 biological replicates). **c**, Western blot of protein collected at 12 hours and 72 hours post mRNA transfection in HEK293T with the constructs indicated in the top row and a *OCT4* promoter targeting gRNA. Anti-Cas9 antibody was used to probe for expression of the construct, while anti-OCT4 antibody was used to probe for protein activation. Anti-tubulin is shown as a loading control. **d**, RT-qPCR analysis of *OCT4* mRNA expression in HEK293T cells 72 hours after transfection with titrating mRNA doses of dCas9-p300 WT or dCas9-p300 I1417N and a constant amount of *OCT4* promoter targeting gRNA (n = 3, mean ± SEM).

We next sought to characterize the transcriptomic impact of expressing these different dCas9-p300 variants by performing RNA-seq on HEK293T cells transiently transfected with specified variants or dCas9 alone and 4 sgRNAs targeting the HS2 enhancer. We found that transfection with dCas9-p300 WT led to substantial perturbation of the transcriptome compared to cells transfected with dCas9 alone when targeting HS2, with 139 differentially expressed genes (DEGs) detected, whereas dCas9-p300 I1417N resulted in 87 DEGs relative to dCas9 alone (**Fig. 3b**). 49 of the DEGs found in dCas9-p300 I1417N were shared between dCas9-p300 I1417N and dCas9-p300 WT transfected cells. Interestingly, most of the observed differentially expressed genes occurred at lowly expressed loci and had small log fold changes, consistent with a model in which the enzymatic activity of exogenously expressed p300 can lead to aberrant H3K27ac enrichment at enhancers and consequently relatively small changes in gene expression in some cases^56,57^. Although global mRNA profiles were similar when comparing transfection with the dCas9-p300 variants against transfection with dCas9 alone, we observed that dCas9-p300 WT displayed the most statistical deviation followed by dCas9-p300 I1417N and then dCas9-p300 D1399Y (**Extended Data Fig. 4d, Supplementary Table 5**). These findings illustrate that although dCas9-p300 I1417N maintains catalytic and transcriptional activity at gRNA-targeted loci, the molecule is less perturbative to the human proteome and transcriptome than dCas9-p300 WT.

### The I1417N mutation destabilizes the dCas9-p300 fusion protein

Because we observed no reduction in enzymatic activity for dCas9-p300 I1417N relative to dCas9-p300 WT (**Fig. 2c,d,e**, **Extended Data 2e**), we hypothesized that the less perturbative proteomic and transcriptomic profiles could be due to the I1417N mutation destabilizing the p300 core domain. Analysis using MUpro^58^, a prediction software for the effect of single-site amino acid mutations on protein stability, showed that the I1417N mutation was predicted to destabilize the p300 core domain protein (ΔΔG = -1.64). To assay the temporal degradation dynamics between different dCas9-p300 variants, we co-transfected mRNAs encoding specific dCas9-p300 variants and a single *OCT4* promoter targeting gRNA in HEK293T cells. mRNA-based delivery of epi-editors has recently emerged as a platform that enables a short burst of effector protein expression prior to mRNA degradation, with effector protein typically absent ∼72 hours after transfection^59^. We collected protein and performed western blotting using anti-Cas9, anti-OCT4, and anti-tubulin antibodies at 9 timepoints spanning 0 to 72 hours post-transfection (**Extended Data Fig. 4e**). Strikingly, when we compared protein levels at 12 and 72 hours, we found that dCas9-p300 I1417N was substantially less abundant than dCas9-p300 WT at both timepoints (**Fig. 3c**). In fact, dCas9-p300 WT protein appeared to be comparable to, if not more abundant than, dCas9-p300 D1399Y at both the 12 and 72 hour timepoints. We observed that OCT4 protein was detectable at ∼12 hours post promoter activation via dCas9-p300 WT and dCas9-p300 I1417N targeting (**Extended Data Fig. 4e**). We additionally titrated the mRNA dose for dCas9-p300 WT and dCas9-p300 I1417N to assess the dose responsivity of both enzymes. We observed that while dCas9-p300 I1417N appeared relatively dose responsive, dCas9-p300 WT saturated gene activation down to the lowest tested dose (15ng; **Fig. 3d**). This result was consistent with our western blot findings (**Fig. 3c, Extended Data Fig. 4e**) and altogether these data indicate that dCas9-p300 I1417N degrades relatively quickly compared to dCas9-p300 WT. Thus, the decreased perturbative and cytotoxic effects of dCas9-p300 I1417N likely arise from the I1417N mutation destabilizing the p300 core domain to a degree that high protein turnover limits aberrant proteomic or transcriptomic perturbations, yet permits on-target acetylation and gene activation.

### dCas9-p300 I1395G catalyzes histone lysine crotonylation and activates transcription when targeted to human promoters

Interestingly, the I1395G mutation previously characterized to skew p300 away from lysine acetylation and towards crotonylation^43^ outperformed rTetR-p300 WT in our rTetR-TetO system (**Fig. 1c**). Crotonylation of histone lysine residues occurs across all four canonical histone subunit proteins as well as on the histone H1 linker protein and is a critical component of signal dependent gene regulation and developmental processes in mammals^36^. For instance, the exogenous addition of crotonate can increase transcription at both reporter genes and endogenous loci^36,60^. Consistent with these studies, we found that histone crotonylation closely correlated with other signatures associated with active transcription when using H3 lysine 18 crotonylation (H3K18cr) from publicly available ChIP-seq data from HCT116 cells^34,61^ as a proxy (**Fig. 4a**). As our initial screening indicated that dCas9-p300 I1395G was functionally active, we next sought to determine if dCas9-p300 I1395G was capable of selectively writing lysine crotonylation.

**Fig 4:**
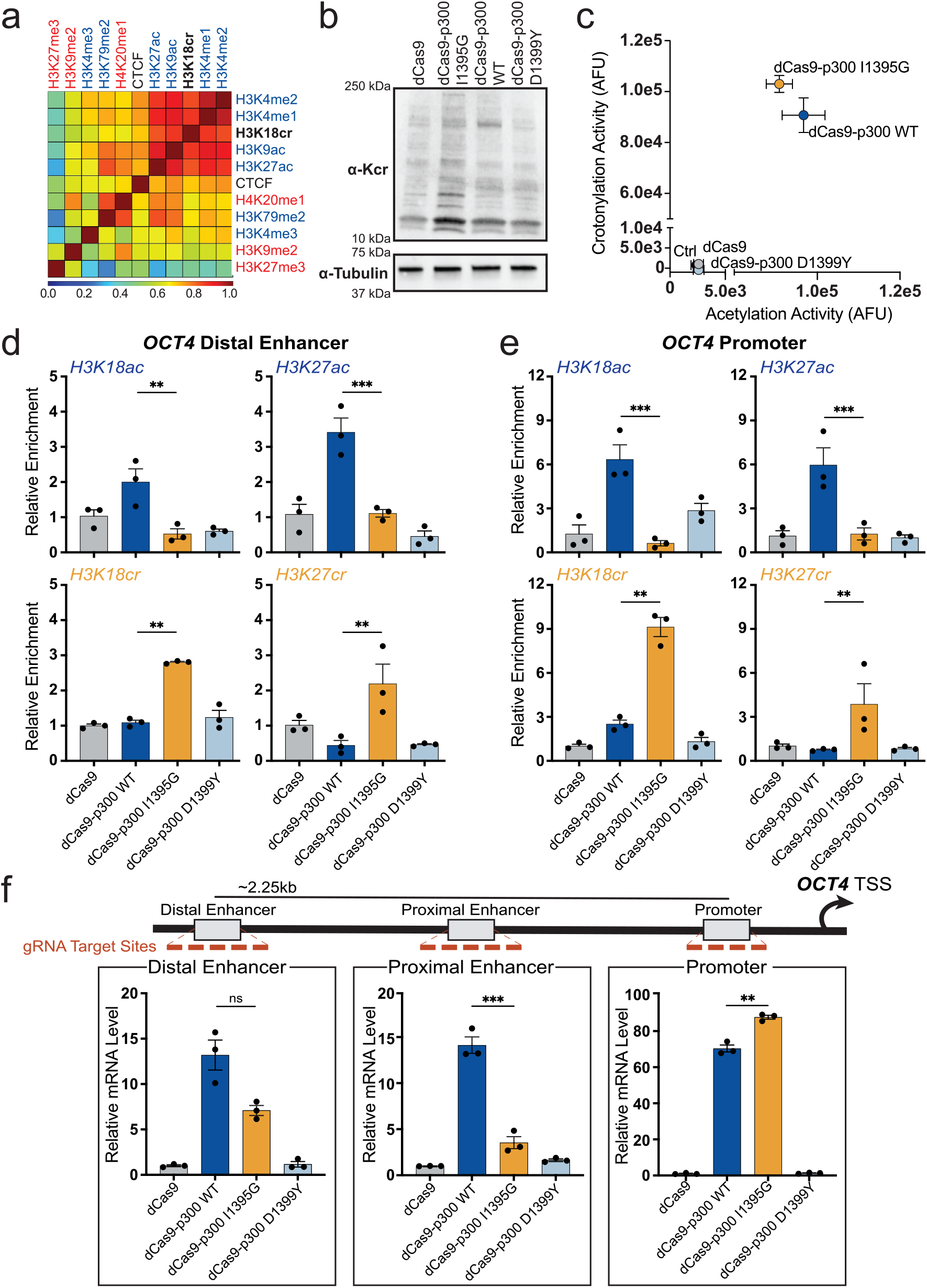
dCas9-p300 I1395G writes lysine crotonylation and activates endogenous human promoters. **a**, Spearman correlation matrix for ChIP-Seq data illustrating the correlation between specific histone modifications or CTCF at specific sites in HCT116 cells. Histone PTM labels in blue are associated with active or poised chromatin states. Histone PTM labels in red are associated with closed or repressive chromatin states. Data collected from GSE96035 for H3K18cr and ENCBS847SOB for remaining groups. **b**, Western blot using an anti-Lysine Crotonylation antibody with protein from HEK293T cells transiently transfected with indicated p300 variants or dCas9 alone. **c**, *In-vitro* acylation activity assay for dCas9-p300 variants (I1395G, D1399Y, WT) and dCas9, along with a no purified protein control (Ctrl), with either Crotonyl-CoA (Y-axis) or Acetyl-CoA (X-axis) co-factor. Activity was measured using the N-terminal H3 peptide (n = 5, mean ± SEM). **d**, Relative enrichment of H3K18/27ac (top, CUT&RUN-qPCR) or H3K18/27cr (bottom, ChiP-qPCR) with indicated p300 variants at the *OCT4* distal enhancer or **e**, the *OCT4* promoter 60 hours after transient transfection with the indicated constructs and respective gRNA pools in HEK293T cells. Relative enrichment was calculated as fold change over dCas9 (n = 3, mean ± SEM). **f**, Top: Schematic of the *OCT4* locus with enhancer and promoter elements arranged relative to the transcription start site (TSS). Red rectangles indicate target gRNA locations. Bottom: RT-qPCR of *OCT4* mRNA expression upon dCas9-p300 variant targeting at the *OCT4* distal enhancer, proximal enhancer, or promoter in HEK293T cells (n = 3, mean ± SEM). Comparison lines throughout indicate multiple comparison testing with Šidák, Dunnett’s T3, or Dunn’s multiple comparisons tests depending on normality, tested via Shapiro-Wilk (*P>0*.*05*), and homoscedasticity, tested via Brown-Forsythe (*P>0*.*05*). ^*^*P*≤*0*.*05*, ^**^*P*≤*0*.*01*, ^***^*P*≤*0*.*001*, ^****^*P*≤*0*.*0001*, ns, not significant.

Upon overexpression of dCas9-p300 I1395G in HEK293T cells, we observed a marked increase of crotonylated lysine residues across the proteome in comparison to dCas9-p300 WT or dCas9-p300 D1399Y (**Fig. 4b**). In addition, we found that *in-vitro* purified dCas9-p300 I1395G crotonylated histone H3 peptides to a greater degree than purified dCas9-p300 WT, with decreased amounts of H3 peptide acetylation observed for the I1395G mutant in a saturating reaction (**Fig. 4c**). To determine the lysine acylation profile of dCas9-p300 I1395G in cells at endogenous loci, we targeted the distal enhancer and promoter of *OCT4* with gRNA pools for each site and performed CUT&RUN-qPCR (for acetyl marks) or ChIP-qPCR (for crotonyl marks). As expected, we found enrichment of H3K18ac and H3K27ac when dCas9-p300 WT was targeted to either the *OCT4* promoter or the *OCT4* distal enhancer (**Fig. 4d,e**). However, we observed a substantial reduction in enrichment of acetylation with dCas9-p300 I1395G to a level indistinguishable from dCas9 alone or to dCas9-p300 D1399Y. Coincident with this reduction in acetylation, we found that dCas9-p300 I1395G deposited both H3K18cr and H3K27cr whereas dCas9-p300 WT was devoid of detectable crotonylation activity (**Figs. 4d,e**).

We next sought to understand the targeted transcriptional effects of histone crotonylation using pools of gRNAs targeting the *OCT4* distal enhancer, proximal enhancer, or promoter in HEK293T cells. We observed that dCas9-p300 I1395G activated transcription at the *OCT4* promoter to a level that was comparable to dCas9-p300 WT (**Fig. 4f**). Promoter activation using dCas9-p300 I1395G was further confirmed at several different human promoters (**Extended Data Fig. 5a**), in an alternative cell type (**Extended Data Fig. 5b**), and at the protein level for *OCT4* and an alternative target (*CD34*; **Extended Data Figs. 5c,d**). However, in contrast to dCas9-p300 WT, we found that dCas9-p300 I1395G only weakly activated the transcription of *OCT4* when targeted to either the proximal or distal enhancer (**Fig. 4f**). Similar effects were observed at several other endogenous enhancers (**Extended Data Fig. 5e**).

These results establish dCas9-p300 I1395G as an effective tool for depositing histone crotonylation in the human genome. Our data also indicate that histone crotonylation and acetylation can be functionally divergent at endogenous human enhancers, yet largely redundant with respect to transcriptional activation at promoters. Furthermore, these findings demonstrate that transcription can be activated from human promoters through targeting with dCas9-p300 I1395G in the absence of detectable H3K18ac and/or H3K27ac deposition.

## Discussion

Here we have shown that single amino acid mutations within the acyl-CoA binding domain of p300 can be used to reduce the cytotoxicity of the enzyme when exogenously expressed and reshape the acyl deposition behavior of p300. p300 and its paralog CBP are the only known direct writers of H3K27ac, a histone PTM that demarcates active promoters and enhancers^22^. The ability to write this PTM is thus critical for understanding transcriptional activation and mechanistically dissecting how distal and proximal regulatory elements control transcriptional activity^62,63^. However, exogeneous expression of p300 core domain fusions has been associated with off-target activity and cytotoxicity that limits its application in some settings^20,64,65^. Through targeted mutagenesis, we identified the I1417N mutation that when installed in p300, resulted in ablation of the cytotoxicity associated with exogenous expression of the p300 core domain. Remarkably, mitigation of cytotoxicity was not concurrent with loss of on-target enzymatic activity, with our benchmarking of dCas9-p300 I1417N showing that this variant can acetylate histone lysine residues and activate transcription to levels generally on par with p300 WT. We also found that the I1417N mutation in p300 enabled higher efficiency packaging into lentivirus and transduction, which should allow this enzyme to be more easily deployed in cell types or applications in which viral transduction is necessary.

The p300 I1417N variant was an outlier to the general trend that reductions in cytotoxicity were concurrent with reductions in activity (**Extended Data Fig. 1f**). To understand why, we began by profiling the PPIs for both dCas9-p300 WT and dCas9-p300 I1417N. The p300 enzyme has been shown to acetylate other targets in the proteome beyond histones^23^, leading us to hypothesize that p300 cytotoxicity may result from aberrant acetylation across the proteome when the p300 core domain, which lacks flanking regulatory regions, is expressed. To capture possible instances of aberrant acetylation, we found the union set of known protein acetylation targets of full-length p300 catalogued in the p300 acetylome with p300 PPIs captured with our exogenously expressed core domain variants^23^. Remarkably, these comparisons demonstrated that p300 WT resulted in over 3 times the number of aberrant acetylation events compared to p300 I1417N. This result, along with global increases in acetylated lysine residues observed with dCas9-p300 WT but not dCas9-p300 I1417N^20^, points towards aberrant protein acetylation as a key driver of p300 associated cytotoxicity. This conclusion is further supported by the observation that although p300 I1417N expression resulted in less perturbations to the human transcriptome than p300 WT, the difference in transcriptomic impact was minimal compared to differences in PPIs. Further, the differentially expressed genes that were observed in our studies were generally lowly expressed genes with minimal fold change over dCas9 alone.

Assaying the degradation dynamics of our dCas9-p300 variants provided further insight into why p300 I1417N was less perturbative than p300 WT. Specifically, the I1417N mutation appeared to destabilize the p300 core domain protein, resulting in dCas9-p300 I1417N being undetectable by western blot within 72 hours after transfection as mRNA, and being dose-responsive to mRNA titration whereas dCas9-p300 WT was not. This was unexpected as we observed that no significant loss in enzymatic activity was incurred due to the I1417N mutation. Thus, we conclude that the I1417N mutation results in a p300 core domain structure that sits in a goldilocks zone of protein stability, whereby on-target histone acetylation and gene activation is still possible, but the protein is degraded before a significant number of off-target perturbations can be incurred. In this model, when expressing dCas9-p300 I1417N via mRNA that only produces a short burst of protein expression, the target site is likely not able to be saturated resulting in the reduced activation observed with dCas9-p300 I1417N mRNA compared to dCas9-p300 WT mRNA. However, when dCas9-p300 I1417N is expressed via a plasmid construct or an integrated cassette, the turnover of dCas9-p300 I1417N is balanced by continual expression resulting in comparable levels of histone acetylation and gene activation when compared with dCas9-p300 WT, while incurring fewer off-target perturbations due to each individual dCas9-p300 I1417N molecule’s faster relative degradation.

In addition to developing the less perturbative dCas9-p300 I1417N programmable acetyltransferase, we also built a programmable crotonyltransferase using the p300 I1395G mutation^43^. We established that dCas9-p300 I1395G can deposit histone crotonylation at specified sites in the human genome, and that histone crotonylation can in turn activate gene transcription when targeted to endogenous promoters. Interestingly, this activation was comparable to an acetylation-competent version of the p300 core. Although we demonstrated that dCas9-p300 I1395G can deposit histone crotonylation at both enhancers and promoters, we only observed meaningful transcriptional activation at promoters, suggesting that histone crotonylation alone may not be sufficient to activate genes from enhancers, at least at the enhancers tested here. This is aligned with enhancer mediated gene activation largely being regulated by bromodomain family proteins such as BRD4, many of which are unable to bind crotonylated residues^39,66,67^. Additionally, while it was recently reported that the YEATS-domain containing protein GAS41 can bind to H3K27cr and repress transcription, we did not observe any transcriptional repression at our tested loci^68^, possibly due to context dependencies or competition by transcription-activating YEATS domain-containing proteins such as ENL or AF9^69^. Taken together, our data suggests that dCas9-p300 I1395G deposits crotonylation at targeted promoters and enhancers without the need for global metabolite supplementation, and that crotonylation has the potential to activate transcription via complementary or orthogonal mechanisms to dCas9-p300 WT, such as YEATS domain mediated gene activation at promoters in tissues where crotonyl-CoA is more prevalent^32,34^.

Overall, our study arms the community with new tools to probe and manipulate endogenous histone acylation in a manner that reduces cytotoxicity and unintended perturbations. These advances are important for biological discovery, human cell engineering, and in the longer term the clinical application of epi-editors. Our data also suggest that methods aimed at reducing the half-life or expression of epi-editing effector domains^65^ through targeted mutagenesis or other means is viable strategy for minimizing off-target effects while preserving on-target activity. While our study here focused on the p300 enzyme due to its unique capabilities and utility, it is likely that other epi-editing effector domains^70^ will be amenable to similar data-driven optimization using either targeted or high-throughput mutagenesis campaigns.

## Methods

### Plasmid construction

For SpdCas9 encoding vectors (Addgene #52961), cloning backbones were modified to have C-terminal entry sites for insertion of different effector domains. The p300 core domain was amplified from pLV-dCas9-p300-P2A-PuroR (Addgene #83889). Mutations within p300 were installed by PCR amplifying fragments of p300 with the variants encoded within and assembled into backbones via NEBuilder HiFi DNA Assembly (NEB, #E2621). Similar strategies were employed for other backbones if C-terminal entry sites did not already exist, including ddCas12a (Addgene #128136), dCasMINI (Addgene #176269), and lenti-pEF-rTetR(SE-G72P)-3XFLAG-LibCloneSite-T2A-mCherry-BSD-WPRE (Addgene #161926). To generate the entry vector for scanning mutagenesis library cloning, p300 was shuttled into an intermediate vector (Addgene #79770), linearized by PCR at the site of interest, and an oligo encoding Esp3I sites was inserted in NEBuilder HiFi DNA Assembly. To generate the reporter plasmid with an SV40 core promoter, subcloning was performed to insert a gBlock (IDT) in the parental reporter vector, AAVS1-PuroR-9xTetO-minCMV-IGKleader-hIgG1_FC-Myc-PDGFRb-T2A-Citrine-PolyA (Addgene #161928). Protein sequences of all dCas9 constructs are shown in **Supplementary Note 1**.

### gRNA cloning

CRISPR gRNAs designed in this study were created using the CRISPR gRNA design tool from Benchling. Oligonucleotides encoding protospacer sequences were ordered from IDT, and cloned as described previously^11^. Briefly, gRNA cloning backbone (SpdCas9 – Addgene, #47108, ddCas12a – Addgene, #128136, dCasMINI – Addgene, #180280, SpdCas9 for scCRISPRa experiment – Addgene, #192506) was digested with corresponding enzymes (Esp3I, NEB, #R0734S; SapI, NEB, #R0569S or BbsI, NEB, #R3539S). Oligos were annealed, phosphorylated, and ligated into the cloning vector using T4 Ligase (NEB, #M0202L). All gRNA protospacer sequences used in this study for SpdCas9, dCas912a, and dCasMINI are listed in **Supplementary Table 6**.

### Cell culture and transfection

HEK293T (ATCC, CRL-11268), HeLa (ATCC, CCL-2), U2OS (ATCC, HTB-96), K562 (ATCC, CRL-243), and MSC (ATCC, PCS-500-011) cells were purchased from American Type Cell Culture (ATCC, USA) and cultured in ATCC-recommended media supplemented with 10% FBS (Sigma-Aldrich) and 1% pen/strep (100units/ mL penicillin, 100μg/ mL streptomycin; Gibco) at 37° C and 5% CO2. MSCs were cultured in ATCC MSC Basal Medium supplemented with MSC Growth Kit (PCS-500-040). All experiments were performed within 20 passages of cell stock thaws, or 10 passages for MSCs. Trypsinization was performed for cells through incubation with Gibco Trypsin-EDTA 0.25% (#25200056) at 37C. All transfections were carried out using either Lipofectamine 3000 (plasmid) or Lipofectamine MessengerMax (mRNA) according to the manufacturer’s instructions.

### Western blotting

Cells were trypsinized at indicated times post-transfection and spun down. Cells were then lysed in RIPA buffer (Thermo Scientific, #89900) with 1X protease inhibitor cocktail (Thermo Scientific, #78442). Lysates were cleared by centrifugation, and total protein quantitation was performed using the BCA method (Pierce, #23225). 15-50μg of lysate were separated using precast 7.5%, 10%, or 4–20% SDS-PAGE (Bio-Rad) and then transferred onto PVDF membranes using the Transblot-turbo system (Bio-Rad). Membranes were blocked with EveryBlot Blocking Buffer (Bio-Rad, #12010020) and incubated overnight at 4°C with primary antibody (anti-Cas9; 1:1000 dilution, Diagenode #C15200216, anti-FLAG; 1:2000 dilution, Sigma-Aldrich #F1804, anti-OCT4, 1:1000 dilution, Cell Signaling technology, #9656, anti-β-Tubulin; 1:1000 dilution, Bio-Rad, #12004166). The membranes were washed with 1X TBST 3 times (5 mins each wash) and incubated with respective HRP-tagged secondary antibodies (1:2000 dilution) for 1 hour at room temperature. Next membranes were washed with 1X TBST 3 times (5 mins each wash. Membranes were then incubated with ECL solution (BioRad, #1705061) and imaged using a Chemidoc-MP system (BioRad). The β-Tubulin antibody was tagged with Rhodamine (Bio-Rad, #12004166) and was imaged using Rhodamine channel in Chemidoc-MP as per manufacturer’s instruction.

### Reverse-transcription quantitative PCR (RT-qPCR)

Cells were trypsinized at indicated times post-transfection and spun down. RNA (including pre-miRNA) was isolated using the RNeasy Plus mini kit (Qiagen, #74136). 500-2000ng of RNA (quantified using Nanodrop 3000C; ThermoFisher) was used as a template for cDNA synthesis (Bio-Rad, #1725038). cDNA was diluted 10X and 4.5 μL of diluted cDNA was used for each qPCR reaction in a 10ul reaction volume. Real-time quantitative PCR was performed using SYBR Green Master Mix (Bio-Rad, #1725275) in the CFX96 Real-Time PCR system with a C1000 Thermal Cycler (Bio-Rad). Results are represented as fold change above control after normalization to *GAPDH* in all experiments using human cells. For murine cells, 18s rRNA was used for normalization. Undetectable samples were assigned a Ct value of 45 cycles. All qPCR primers and cycling conditions are listed in **Supplementary Table 6**.

### Flow-cytometry

HEK293T rTetR-rTetO reporter cells, or HEK293T cells, were collected 72 hours post-transduction/transfection via trypsinization. Single-cell suspensions were washed with complete media and then with 0.22 μM filtered 1 × FACS buffer (1% BSA in 1 × PBS). For CD34 expression analysis, cells were incubated with CD34-PE antibody (20 μL per 10^6^ cells) or IgG-PE isotype antibody (Invitrogen, no. 12-4714-42) in 1 × FACS buffer for 30 min. HEK293T rTetR-rTetO reporter cells were directly run on the flow cytometer. Stained cell fluorescence intensity was measured using a Sony SA3800 spectral analyzer. Data were analyzed using FlowJo software (v.10). First, cell debris and doublets were excluded with the FSC-A/SSC-A dot plot, followed by the FSC-H/FSC-A dot plot. Negative CD34 gates were set using unstained controls. Negative mCherry and negative Citrine gates were set using untransduced controls. Representative gating strategy shown in **Supplementary Fig. 2**.

### Lentivirus production and titering

One day before transfection, HEK293T cells were seeded at ∼40% confluency in a 10cm^2^ plate. The next day cells were transfected at ∼80–90% confluency. For each transfection, 10μg of plasmid containing the vector of interest, 10μg of pMD2.G (Addgene, #12259), and 15μg of psPAX2 (Addgene, #12260) were transfected using Lipofectamine 3000 (Invitrogen, L3000015) or calcium phosphate. 14 hours post-transfection the media was changed. Supernatant was harvested 24 and 48 h post-transfection and filtered with a 0.45μm PVDF filter (Millipore, #SLGVM33RS). Virus was then concentrated at 100X using Lenti-X™ Concentrator (Takara, 631232), aliquoted and stored at −80 °C. Lentiviral titers were measured by the Lenti-X™ qRT-PCR Titration Kit (Takara, #631232).

### In-vitro acylation activity assay

Wild-type or I1395G dCas9-p300 were expressed and purified from transiently transfected HEK293T cells using Anti-FLAG M2 Affinity Gel (Millipore Sigma, A2220). In vitro HAT and HCT assays were performed according to manufacturer’s instructions (Active Motif, 56100). Briefly, 1ug of recombinant protein was incubated in Assay Buffer AM1 with 125uM of either acetyl-CoA (Active Motif, 56100) or crotonyl-CoA (Sigma, 28007) and 50uM of Histone H3 peptide for 30 min at room temperature. The reaction was terminated with Stop Solution, incubated with Developing Solution for 15 min at room temperature protected from light, and quantified via fluorescence with excitation at 360-390nm and emission at 450-470nm.

### ChIP-qPCR

HEK293T cells were co-transfected with indicated dCas9 fusion (7.5 μg) expression vectors and gRNA (2.5 μg) constructs in 10 cm plates in biological duplicates for each condition tested. ∼72 hours post-transfection, cells were cross-linked for 10 min at RT using 1% formaldehyde (Sigma-Aldrich, F8775-25ML) and then the reaction was stopped by the addition of glycine to a final concentration of 125 mM. Cells were harvested and washed with ice cold 1X PBS and suspended in Farnham lysis buffer (5 mM PIPES pH 8.0, 85 mM KCl, 0.5% NP-40) supplemented with protease inhibitor (Thermo Scientific, A32965). Cells were then pelleted and resuspended in RIPA buffer (1X PBS, 1% NP-40, 0.5% sodium deoxycholate, 0.1% SDS) supplemented with protease inhibitor. Approximately 2.5e7 cells were used for each ChIP experiment. Chromatin in RIPA buffer was sheared to a median fragment size of around 250 bp using a Bioruptor XL (Diagenode). 5 μg of α-H3K27Cr (Thermo Fisher, #712478) or α-H3K18Cr antibody (Thermo Fisher, #703472) was incubated with 50 μl Rabbit IgG magnetic beads (Life Technologies, 11203D) for ∼4 hours at 4 °C, respectively. Antibody-linked magnetic beads were washed 3 times with PBS/BSA buffer (1X PBS and 5 mg/ml BSA) and sheared chromatin was incubated with corresponding antibody-linked magnetic beads at 4° C overnight and then washed 5 times with LiCl IP wash buffer (100 mM Tris pH 7.5, 500 mM LiCl, 1% NP-40, 1% sodium deoxycholate). Cross-links were then reversed via overnight incubation at 65°C and DNA was purified using QIAquick PCR purification kit (Qiagen, 28106) for ChIP-seq. Input DNA was prepared from ∼1.0e6 cells. 10 ng of DNA was used for subsequent qPCR reactions using a CFX96 Real-Time PCR Detection System with a C1000 Thermal Cycler (Bio-Rad, #1855195). Baselines were subtracted using the baseline subtraction curve fit analysis mode and thresholds were automatically calculated using the Bio-Rad CFX Manager software version 2.1. The ChIP-qPCR data was relative to percent input. All ChIP-qPCR primers and conditions are listed in **Supplementary Table 6**.

### CUT&RUN

CUT&RUN was performed using the EpiCypher CUTANA ChIC/CUT&RUN Kit (EpiCypher, #14-1048). Briefly, 500k HEK293T cells were detached and harvested using 0.5mM EDTA (Fisher, #BP2482-500), washed once with 1X PBS and then resuspended in 300 μl of wash buffer. Next each of 3 100 μl aliquots (∼1/3 of each 24 well) of cells were processed for H3K4me3 antibody (EpiCypher, #13-0041), H3K27ac antibody (EpiCypher, #13-0045), H3K27me3 antibody (EpiCypher, #13-0055), H2BK20ac antibody (Abcam, ab177430), or input DNA, respectively. Cells were first immobilized on concanavalin A beads, and then incubated with respective antibody (0.5ug/sample) overnight at 4°C in antibody dilution buffer (cell permeabilization buffer + EDTA). On the following day, cells were washed twice with cell permeabilization buffer. After washing the beads, pAG-MNase was added to the immobilized cells and then incubated for 2 hours at 4°C to digest and release DNA. Libraries were prepared using the CUT&RUN Library Prep Kit (EpiCypher, #14-1001) and DNA was quantified using the Qubit HS dsDNA High Sensitivity Kit (Thermo Fisher) and sequenced at a depth of 15M paired end reads per sample on a NextSeq 2000 (High Output 2×100). Samples were normalized using an E. Coli spike-in DNA (EpiCypher, #18-1401). For CUT&RUN-qPCR assays, purified DNA from either H3K4me3, H3K27ac, H3K27me3, or H2BK20ac antibody incubated samples were then assayed by qPCR. Relative enrichment of H3K4me3 and H3K27ac was expressed as fold change above control cells transfected with dCas9 plasmid and after normalization to purified input DNA. qPCR primers used for CUT&RUN are shown in **Supplementary Table 6**.

### CUT&RUN sequencing analysis

Data was analyzed using CUTANA Cloud by EpiCypher using CUTANA™ CUT&RUN/Tag Alignment app version 0.0.12. FASTQ files were aligned to Human hg38 using Bowtie2 (version 2.2.9). SAMtools (version 1.13) was used to remove multi-aligned reads. BEDTools (version 2.30.0) was used to remove reads aligned to the ENCODE DAC Exclusion List regions. Duplicate reads were filtered out using Picard (version 2.27.1). RPKM-normalized bigWigs were generated using deepTools (version 3.5.1) with –binsize set to 20. The computeMatrix utility from deepTools was used to calculate target enrichment relative to transcription start sites (reference-point mode) and annotated gene bodies (scale-regions mode), and heatmaps were generated with the plotheatmap utility from deepTools. Smoothed Bigwigs were generated for visualization in the Integrative Genomics Viewer (IGV), using the –smoothLength option set to 100.

### Fluorescent imaging

HEK293T cells were fixed with 4% formaldehyde (Millipore Sigma, #47608) in PBS (Genesee, 25-508), then permeabilized with 0.5% Triton^®^ X-100 in PBS. After permeabilization, HEK293T cells were stained with 1:1000 DAPI (Thermo Fisher, 62247), then imaged on an Eclipse Ti2-E inverted microscope 48 hours transfection. All samples from each experiment were imaged with the same excitation intensity and exposed at the same exposure time.

### Cell viability assay

A 1:10 dilution of alamarBlue (Invitrogen, DAL1025) was directly added to wells with cultured MSCs or K562 cells, and incubated at 37°C for 2 hours. After 2 hours, fluorescence was read with an excitation emission of 540 nm and an emission wavelength of 580 nm on the BMG CLARIOstar plate reader. Relative viability was calculated as the ratio of each samples fluorescence emission to that of a control sample transfected with dCas9 alone.

### Immunoprecipitation mass spectrometry

Cells were harvested with a cell scraper and spun at 300xg for 5 min. The packed cell volume was measured and 2.5 volumes of NETN buffer was added to the cell pellet. (NETN: 50mM Tris pH 7.3, 170mM NaCl, 1mM EDTA, 0.5% NP-40). Lysate was sonicated (30sec bust, 59sec off, repeat 6 times) using a Diagenode Bioruptor Pico (#B01060010) and then centrifuged at 20,000xg, for 20min at 4°C. Supernatant was harvested and then transferred to a fresh tube. 5µg of FLAG antibody (Sigma-Aldrich, #F1804-200UG) was added to the above supernatant and incubated for 1 hour at 4°C rocking. Supernatant was spun again at 20,000xg for 20min and the supernatant was collected in a fresh tube. 20 µl of protein A bead slurry was added to the above supernatant and incubated for 1 hour at 4°C with inverted rotation. Supernatant and beads were spun at 1000xg for 1min. Flow-through was saved for downstream QC testing. Beads were washed with 1ml NETN buffer and spun at 1000xg for 1min and supernatant was discarded. A flat-ended tip was used to remove residual buffer solution from the beads completely. 20µl of 2X SDS loading dye was added to the beads and samples were heated for 8-10 min at 90°C. The immunoprecipitated samples were resolved on NuPAGE 10% Bis-Tris Gel (Life Technologies), each lane was excised into 6 equal pieces and combined into two peptide pools after in-gel digestion using trypsin enzyme. The peptides were dried in a speed vac and dissolved in 5% methanol containing 0.1% formic acid buffer. The LC-MS/MS analysis was carried out using the nano-LC 1000 system coupled to Orbitrap Fusion mass spectrometer (Thermo Scientific). Peptides were eluted on an analytical column (20 cm x 75 μm I.D.) filled with Reprosil-Pur Basic C18 (1.9 µm, Dr. Maisch GmbH, Germany) using 110 ml discontinuous gradient of 90% acetonitrile buffer (B) in 0.1% formic acid at 200 nl/min (2-30% B: 86 min, 30-60% B: 6 min, 60-90% B: 8 min, 90-50% B: 10 min). The full MS scan was performed in Orbitrap analyzer in the range of 300-1400m/z at 120,000 resolution followed by an IonTrap HCD-MS2 fragmentation for a cycle time of 3 seconds with precursor isolation window of 3m/z, collision energy 30%, AGC of 50000, maximum injection time of 30 ms.

The MS raw data were searched using Proteome Discoverer 2.0 software (Thermo Scientific) with Mascot algorithm against human NCBI RefSeq database updated 2020_0324. The precursor ion tolerance and product ion tolerance were set to 20 ppm and 0.5 Da, respectively. Maximum cleavage of 2 with Trypsin enzyme, dynamic modification of oxidation (M), protein N-term acetylation, deamidation (N/Q) and destreak (C) was allowed. The peptides identified from the mascot result file were validated with a 5% false discovery rate (FDR). The gene product inference and quantification were done with label-free iBAQ approach using the ‘gpGrouper’ algorithm. For statistical assessment, missing value imputation was employed through sampling a normal distribution N (μ-1.8 σ, 0.8σ), where μ, σ are the mean and standard deviation of the quantified values. For differential analysis, we used the moderated t-test and log2 fold changes as implemented in the R package limma^71^ and multiple-hypothesis testing correction was performed with the Benjamini–Hochberg procedure. To filter out background contaminants, we removed from further analysis proteins that were recovered in more than 50% of IP analyses in HEK293T cells according to the contaminant repository for affinity purification (CRAPome^52^). To quantify shared hits in the STRING database, all interactors of p300 with a combined score > 0.15 were extracted and compared against hits found in this study’s IP-MS dataset.

### RNA-Sequencing

RNA sequencing (RNA-seq) was performed in duplicate for each experimental condition. RNA was isolated from transfected cells using the RNeasy Plus mini kit (Qiagen, #74136). RNA-seq libraries were constructed using the TruSeq Stranded Total RNA Gold (Illumina, #RS-122-2303). The qualities of RNA-seq libraries were verified using the Tape Station D1000 assay (Tape Station 2200, Agilent Technologies), and the quantities of RNA-seq libraries were checked again using real-time PCR (QuantStudio 6 Flex Real time PCR System, Applied Biosystem). Libraries were prepared with ERCC92 spike-in controls, normalized, pooled, and then 75 bp paired-end reads were sequenced on the HiSeq3000 platform (Illumina). Adapter trimming and quality filtering were performed using *fastp* (v0.23.2), with quality control metrics recorded in HTML and JSON formats. Reads were aligned to a custom reference genome combining the human genome (hg38) and ERCC92 spike-in sequences using *HISAT2* (v2.2.1). The reference was built by merging *hg38*.*fa* and *ERCC92*.*fa*, along with their respective annotation files (*hg38*.*gtf* and *ERCC92*.*gtf*). Paired-end, strand-specific alignments were then performed and SAM files were generated for downstream analysis. Gene expression levels were quantified using *RSEM* (v1.3.3)^72^, processing the aligned reads to simultaneously measure expression of both endogenous genes and ERCC spike-ins^73^. For spike-in and gene-level quantification, *featureCounts* was used to assess read counts, enabling normalization of transcript abundance^74^. Differential expression analysis was performed using DESeq2^75^. Volcano plots, PCA plots, and heat maps were created on DESeq2 using the analyzed data.

### In-Vitro transcription

mRNA was generated using template DNA encoding our dCas9-p300 variants of interest driven off a T7 RNA Polymerase. Briefly, template plasmid was digested with HINDIII-HF (NEB, R3104S). 1 μg of linearized template DNA was then used in the HiScribe^®^ T7 High Yield RNA Synthesis Kit (NEB, E2040S) in-vitro transcription reaction with CleanCap^®^ Reagent AG (TriLink, N-7113) and complete substitution of UTP with N1-Methylpseudouridine-5’-Triphosphate (TriLink, N-1081). Purification of mRNA was done using Lithium Chloride solution (Sigma, L9650). OCT4 gRNA A used for mRNA experiments was ordered directly from Synthego.

### Data Availability

The raw and processed P300 screening data, RNA-seq, CUT&RUN, and scRNA-seq data are publicly available at GEO (GSE255610). The mass spectrometry proteomics data have been deposited to the ProteomeXchange Consortium via the PRIDE partner repository with the dataset identifier PXD050050. Other data supporting the findings of this work are provided within the article and its **Supplementary Information. Source data** are provided with this paper. Plasmids will be deposited to Addgene upon publication.

### Code Availability

No new or custom code was generated as part of this research.

## Acknowledgements

The authors thank all members of the Hilton and Diehl labs for helpful discussions and insights. The authors also thank Drs. Martis Cowles and Andrea Johnstone (EpiCypher) for generously providing reagents and scientific input on the project. This work was supported by a Cancer Prevention & Research Institute of Texas (CPRIT) award (RR170030) and NIH awards (R35GM143532 and R56HG012206) to I.B.H and a Welch Foundation award (C-2240-20250403) to I.B.H. and M.R.D. J.G. was supported by NSF NRT Program 1828869. M.E. was supported by the American Heart Association awards (917025 and https://doi.org/10.58275/AHA.25TPA1463933.pc.gr.233910) and a ZOLL foundation award.

This project was supported in part by the Baylor College of Medicine (BCM) Genomic and RNA Profiling Core with funding from NIH NCI (P30CA125123), NIH S10 grant (1S10OD023469), and CPRIT (RP200504). This project was also supported by the BCM Single Cell Genomics Core with funding from the NIH (S10OD018033, S10OD023469, S10OD025240) and P30EY002520. The project also received support from the BCM Mass Spectrometry Proteomics Core. The BCM Mass Spectrometry Proteomics Core is supported by the Dan L. Duncan Comprehensive Cancer Center NIH award (P30 CA125123) and a CPRIT Core Facility Award (RP210227).

## Contributions

I.B.H., J.G., and R.J., conceptualized the research project and designed the experiments. J.G., R.J., J.L., K.G., G.C.B., D.R., B.M., and S.K., designed and cloned experimental constructs and gRNAs. J.G. and R.J. analyzed the mass spectrometry data. R.J. and A.J.M. analyzed the CUT&RUN and RNA sequencing data. J.G., R.J., and J.L. performed the CUT&RUN and ChIP experiments. J.G. and D.R. performed the flow-cytometry, lentivirus, and *in-vitro* acylation experiments. J.G., R.J., and B.M. performed the western blot experiments. J.G., R.J., and K.G. performed the imaging experiments. J.G., R.J., and D.R. performed the RT-qPCR experiments. R.J. and G.C.B produced mRNA, cultured Mesenchymal Stem cells, and ran the time-course western blot. R.J. performed cell viability experiments. G.C.B., K.E.M., M.R.D., M.E., and I.B.H. provided assistance with experimental designs and conceptual discussions. I.B.H. supervised the work. R.J., J.G., M.E. and I.B.H. wrote the manuscript with inputs all authors.

## Corresponding author

Correspondence to Isaac Hilton

## Ethics declarations

J.G., J.L., B.M., M.E., and I.B.H. are inventors on patents related to genome and epigenome editing. K.E.M is an employee of EpiCypher Inc. The remaining authors declare no competing interests.

---

**Extended Data Fig 1:**
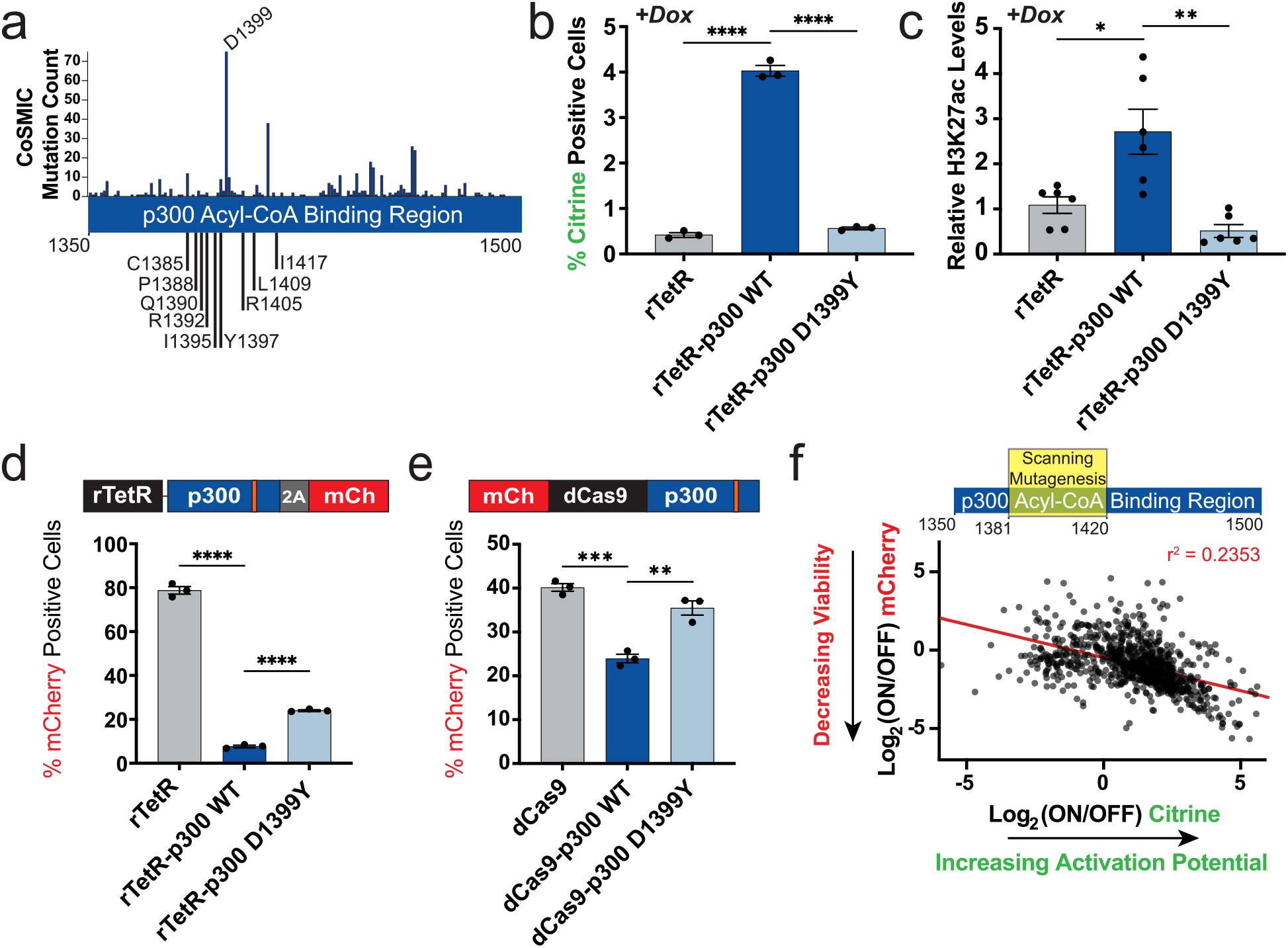
The rTetR-rTetO screening system enables evaluation of p300 variant expression and activation potential. **a**, Catalogue of Somatic Mutations in Cancer (COSMIC) database mutation counts across the p300 Acyl-CoA binding region (1350-1500), with tested p300 variants annotated below. Mutations labeled on the bottom were not found in the COSMIC database. **b**, Citrine flow-cytometry readouts for the rTetR-rTetO screening system transduced with the indicated rTetR constructs and in the presence of doxycycline, 48 hours post doxycycline addition (n = 3, mean ± SEM). **c**, CUT&RUN-qPCR for H3K27ac of the reporter locus in HEK293T cells stably transduced with indicated rTetR-p300 variants 48 hours post-doxycycline addition (n = 6, mean ± SEM). **d**, mCherry flow-cytometry readout for the rTetR-rTetO screening system transduced with the indicated rTetR constructs and in the presence of doxycycline, 48 hours post doxycycline addition (n = 3, mean ± SEM). **e**, mCherry flow-cytometry readout for dCas9-p300 variants directly fused to mCherry, 48 hours after transfection into HEK293T cells (n = 3, mean ± SEM). **d, e**, Orange line in schematic indicates approximate location of p300 variant mutation. **f**, Scatter plot illustrating the relationship between reporter activation (Log_2_(ON/OFF) Citrine, X-axis) and expression (Log_2_(ON/OFF) mCherry, Y-axis) for 840 scanning mutagenesis variants screened with the rTetR-rTetO system. Line of best fit from a simple linear regression is shown in red (r^2^ = 0.2353). Comparison lines throughout indicate multiple comparison testing with Šidák, Dunnett’s T3, or Dunn’s multiple comparisons tests depending on normality, tested via Shapiro-Wilk (*P>0*.*05*), and homoscedasticity, tested via Brown-Forsythe (*P>0*.*05*). ^*^*P*≤*0*.*05*, ^**^*P*≤*0*.*01*, ^***^*P*≤*0*.*001*, ^****^*P*≤*0*.*0001*, ns, not significant.

**Extended Data Fig 2:**
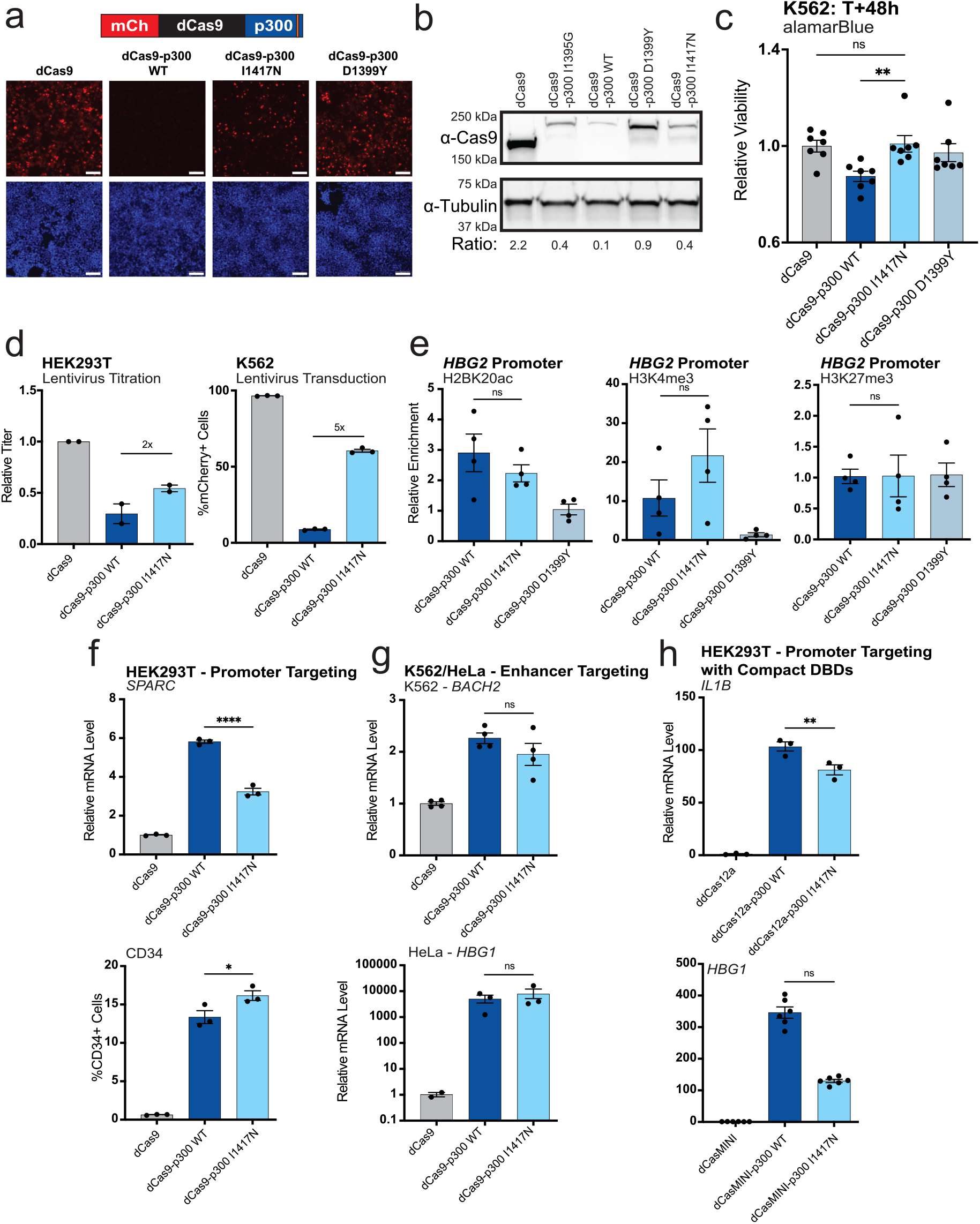
p300 I1417N is broadly compatible with diverse target loci, cell-types, and DNA binding modalities. **a**, Fluorescence microscopy 48 hours after HEK293T cells were transiently transfected with dCas9, dCas9-p300 WT, dCas9-p300 I1417N, or dCas9-p300, all fused N-terminally to a mCherry fluorophore and co-transfected with a scrambled gRNA. Top panels: mCherry channel fluorescence. Bottom panels: DAPI staining. Scale bar: 100um. Orange line in schematic indicates approximate location of p300 variant mutation. **b**, Western blot of protein from HEK293T cells 72 hours after transient transfection with the indicated constructs and a scrambled gRNA using an anti-Cas9 antibody to blot for expression of the indicated construct and an anti-tubulin construct as a loading control. Densitometry ratio of the anti-Cas9 band to the anti-tubulin band shown in the bottom row. **c**, Relative cell viability of K562 cells 48 hours after transfection with the indicated constructs given by relative alamarBlue fluorescence compared to the dCas9 transfected K562s (n = 7, mean ± SEM). **d**, (left) Relative lentivirus tittering results from HEK293T cells transfected with lentiviral packaging plasmids encoding the indicated constructs (with bicistronic expression of mCherry). (right) Transduction efficiency with the indicated lentiviral preps at 2% volume by volume in K562 cells, measured through flow-cytometry assessment of mCherry fluorescence (n = 2-3, mean ± SEM). **e**, CUT&RUN qPCR in HEK293T cells 60 hours after transient transfection with the indicated p300 constructs and *HBG1/2* sgRNAs for (left) H2BK20ac, (middle) H3K4me3, and (right) H3K27me3, with relative enrichment calculated as fold change over the dCas9-p300 D1399Y levels (n = 4, mean ± SEM). **f**, RT-qPCR for (top) *SPARC* 72 hours after transient transfection of the indicated constructs with gRNAs targeted to the respective gene promoters. (bottom) Flow-cytometry analysis of the cell surface marker CD34 in HEK293T cells 72 hours after transient transfection of the indicated constructs with gRNAs targeted to the CD34 promoter (n = 3, mean ± SEM). **g**, RT-qPCR for (top) *BACH2* in K562 cells and (bottom) *HBG1* in HeLa cells 72 hours after transient transfection with the indicated constructs and sgRNAs targeting the gene’s respective enhancers (n = 2-4, mean ± SEM). **h**, RT-qPCR for (top) *IL1B* with ddCas12a constructs and (bottom) *HBG1* with dCasMINI constructs 72 hours after transfection with the indicated constructs and sgRNAs targeting the respective gene’s promoters (n = 3-6, mean ± SEM). Comparison lines throughout indicate multiple comparison testing with Šidák, Dunnett’s T3, or Dunn’s multiple comparisons tests depending on normality, tested via Shapiro-Wilk (*P>0*.*05*), and homoscedasticity, tested via Brown-Forsythe (*P>0*.*05*). ^*^*P*≤*0*.*05*, ^**^*P*≤*0*.*01*, ^***^*P*≤*0*.*001*, ^****^*P*≤*0*.*0001*, ns, not significant.

**Extended Data Fig 3:**
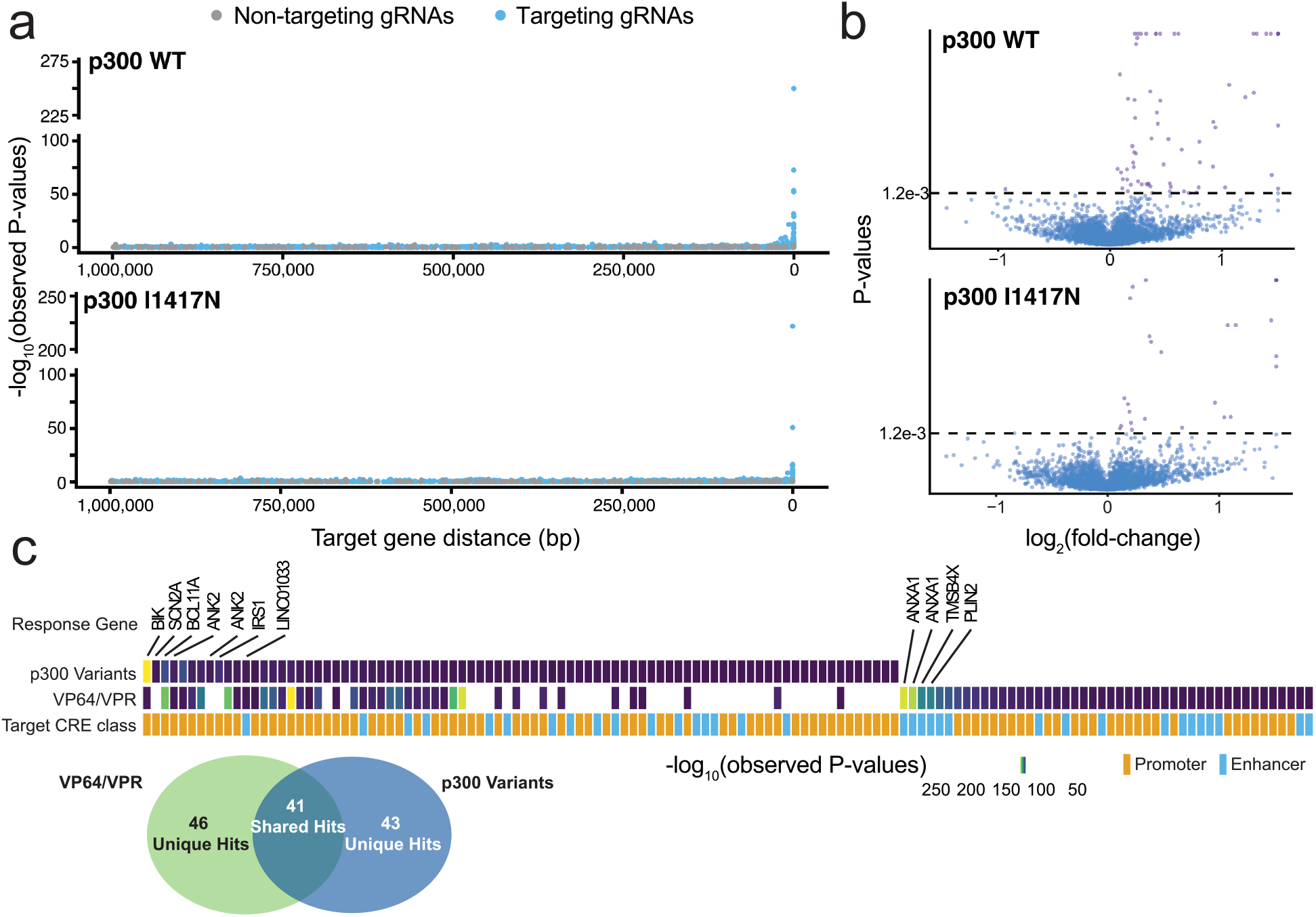
p300 WT and I1417N share perturbation profiles, and uncover unique regulatory elements compared to VP64/VPR associated perturbation. **a**, -log_10_(observed p-values) for differential expression tests from SCEPTRE plotted over the distance from the gRNA target to the tested target gene for (top) dCas9-p300 WT and (bottom) dCas9-p300 I1417N. **b**, Volcano plots showing average log2 fold change and *P*-values for targeting tests for (top) dCas9-p300 WT and (bottom) dCas9-p300 I1417N. Two-tailed SCEPTRE *P*-values throughout were derived using the default SCEPTRE parameters, and subsequently subject to Benjamini-Hochberg correction with those having FDR <0.1 being kept for the two-sided discovery sets. **h**, Perturbations from the p300 variants used in this study were compared against the perturbations detected with VP64/VPR in a previous study (reference 47 in main text). Perturbations were first sorted by decreasing -log_10_(observed *P*-values) for the p300 variants from left to right until the end of the p300 variant perturbations. Remaining perturbations unique to the VP64/VPR dataset were similarly sorted by decreasing -log_10_(observed *P*-values) from left to right. The “Target CRE Class” row was colored depending on if the perturbed element was a promoter or enhancer element. The response gene row contains annotations for perturbations of interest. Venn-Diagram displays unique and shared perturbations identified between the p300 variants and VP64/VPR.

**Extended Data Fig 4:**
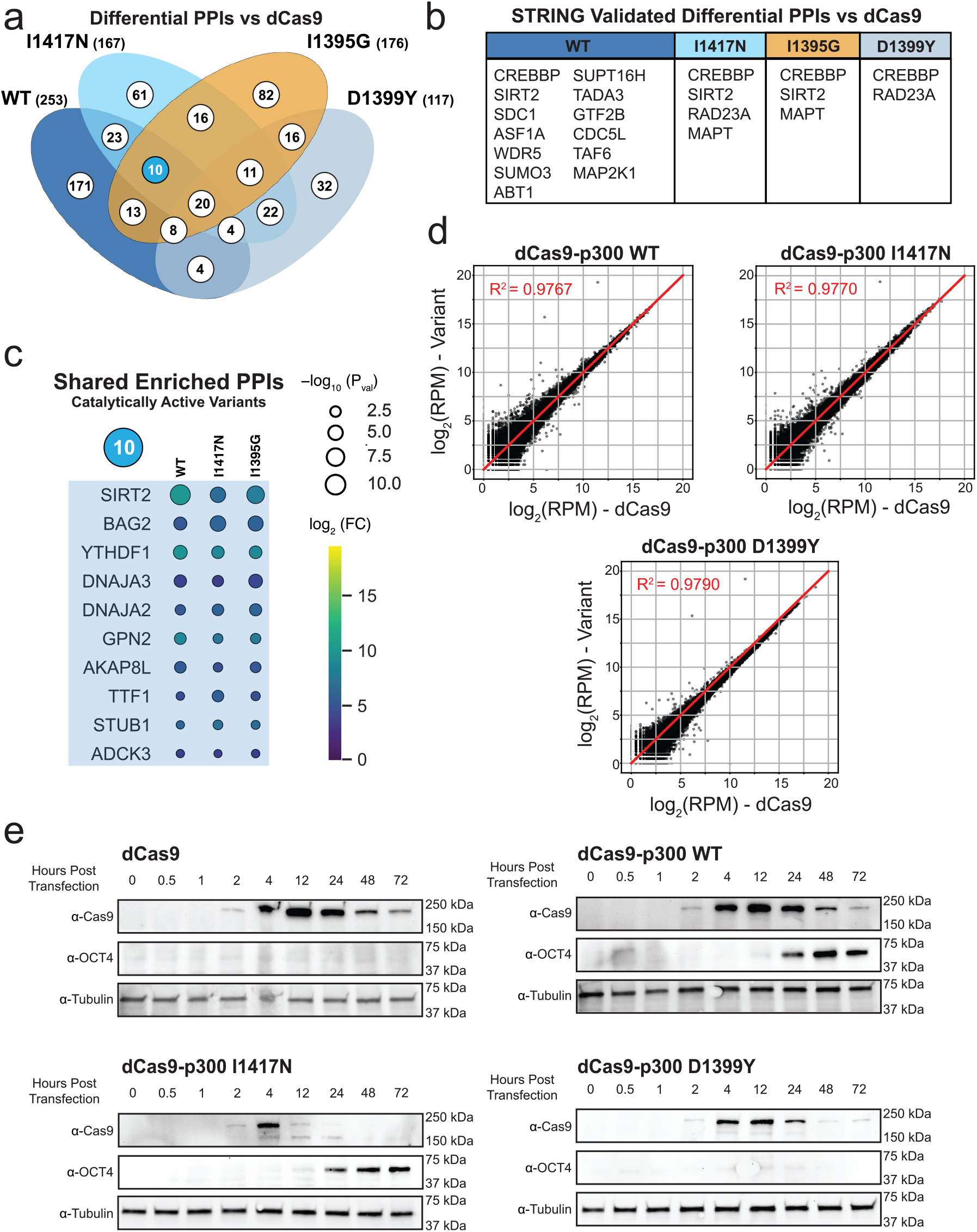
IP-MS, RNAseq, and Western-Blotting for dCas9-p300 variants. **a**, Venn diagram for the four p300 variants used in the IP-MS experiment with values indicating the count of differential protein-protein interactions (PPIs) detected for a given variant or set of variants when compared against a dCas9 only “bait”. Blue colored circle indicates the shared set of PPIs for the catalytically active variants (WT, I1417N, I1395G). **b**, Set of differential PPIs that were also found in the STRING database of known p300 protein interactors for each p300 variant. **c**. Set of differential PPIs shared between the catalytically active p300 variants graphed in order of decreasing average significance from top to bottom. Color of each circle indicates the log_2_(Fold-Change) over the dCas9 “bait” for each protein, and the size of the circle indicates the significance of the differential hit. **d**, log_2_(Reads per Million) of each p300 variant compared against the log_2_(Reads per Million) of dCas9 for all transcripts captured by RNAseq of HEK293T cells transfected with the indicated p300 variants or dCas9 alone and 4 HS2 enhancer element targeting gRNAs (n = 2 biological replicates). Each black dot indicates a single gene and the red line indicates Y=X demarcating perfect correlation between transcriptomes. R^2^ values indicates goodness of fit of the data to Y=X. **e**, Full time course western blots for protein collected at the indicated timepoints from HEK293T cells transfected with mRNA encoding the indicated construct and a single *OCT4* promoter targeting gRNA. Anti-Cas9 antibody was used to assay for expression of the mRNA encoded construct, while anti-OCT4 antibody was used to assay for protein activation. Anti-tubulin is shown as a loading control.

**Extended Data Fig 5:**
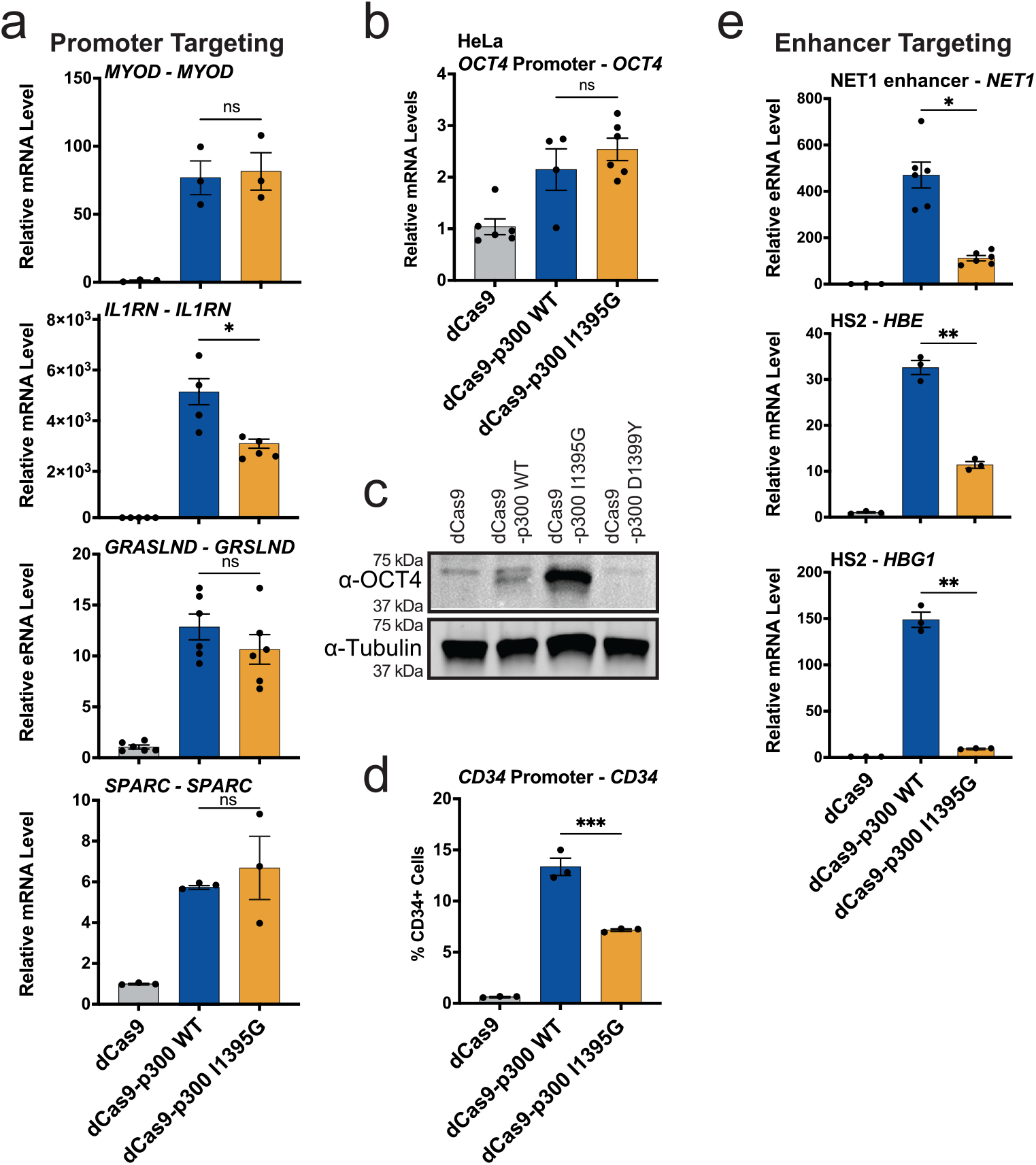
dCas9-p300 I1395G can activate gene transcription from human promoters. **a**, RT-qPCR of *MYOD, IL1RN, GRSLND*, and *SPARC* 72 hours after transient transfection in HEK293T cells of the indicated p300 variants or dCas9 and gRNAs targeted to their respective promoters (n = 3-6, mean ± SEM). **b**, RT-qPCR of *OCT4* 72 hours after transient transfection in HeLa cells of the indicated p300 variants or dCas9 and gRNAs targeting the *OCT4* promoter (n = 4-6, mean ± SEM). **c**, Western blot using an anti-OCT4 antibody on protein collected from HEK293T cells 72 hours post transient transfection of indicated p300 variants or dCas9 and gRNAs targeting the *OCT4* promoter. **d**, Flow cytometry analysis of a cell surface marker CD34 on HEK293Ts 72 hours after transient transfection with the indicated p300 variants or dCas9 and gRNAs targeting the CD34 promoter (n = 3, mean ± SEM). **e**, RT-qPCR of *NET1, HBE*, and *HBG1* 72 hours after transient transfection in HEK293T cells of the indicated p300 variants or dCas9 and gRNAs targeting their respective enhancers (n = 3-6, mean ± SEM). Comparison lines throughout indicate multiple comparison testing with Šidák, Dunnett’s T3, or Dunn’s multiple comparisons tests depending on normality, tested via Shapiro-Wilk (*P>0*.*05*), and homoscedasticity, tested via Brown-Forsythe (*P>0*.*05*). ^*^*P*≤*0*.*05*, ^**^*P*≤*0*.*01*, ^***^*P*≤*0*.*001*, ^****^*P*≤*0*.*0001*, ns, not significant.

